# A catch-bond drives stator mechanosensitivity in the Bacterial Flagellar Motor

**DOI:** 10.1101/174292

**Authors:** AL Nord, E Gachon, R Perez-Carrasco, JA Nirody, A Barducci, RM Berry, F Pedaci

## Abstract

The bacterial flagellar motor (BFM) is the rotary motor which powers the swimming and swarming of many motile bacteria. The torque is provided by stator units, ion motive force powered ion channels known to assemble and disassemble dynamically in the BFM. This turnover is mechano-sensitive, with the number of engaged units dependent upon the viscous load experienced by the motor through the flagellum. However, the molecular mechanism driving BFM mechano-sensitivity is unknown. Here we directly measure the kinetics of arrival and departure of the stator units in individual wild-type motors via analysis of high-resolution recordings of motor speed, while dynamically varying the load on the motor via external magnetic torque. Obtaining the real-time stator stoichiometry before and after periods of forced motor stall, we measure both the number of active stator units at steady-state as a function of the load and the kinetic association and dissociation rates, by fitting the data to a reversible random sequential adsorption model. Our measurements indicate that BFM mechano-sensing relies on the dissociation rate of the stator units, which decreases with increasing load, while their association rate remains constant. This implies that the lifetime of an active stator unit assembled within the BFM increases when a higher force is applied to its anchoring point in the cell wall, providing strong evidence that a catch-bond mechanism can explain the mechano-sensitivity of the BFM.

## 1 Introduction

The bacterial flagellar motor (BFM) is a large molecular complex found in many species of motile bacteria which actively rotates each flagellum of the cell, enabling swimming, chemotaxis, and swarming.^1–5^ The rotor of the BFM is embedded within and spans the cellular membranes, coupling rotation to the extracellular hook and flagellar filament. Multiple transmembrane complexes, called stator units, are anchored around the perimeter of the rotor and bound to the rigid peptidoglycan (PG) layer.^6–8^ In *E. coli*, each stator is composed of MotA and MotB proteins, the latter of which forms an ion channel and allows the stator unit to harness energy from the proton motive force^9–12^ (Fig. 1A). The stator units are responsible for torque generation, acting upon the common track formed by FliG units on the cytosolic side of the rotor.^13, 14^

**Figure 1:**
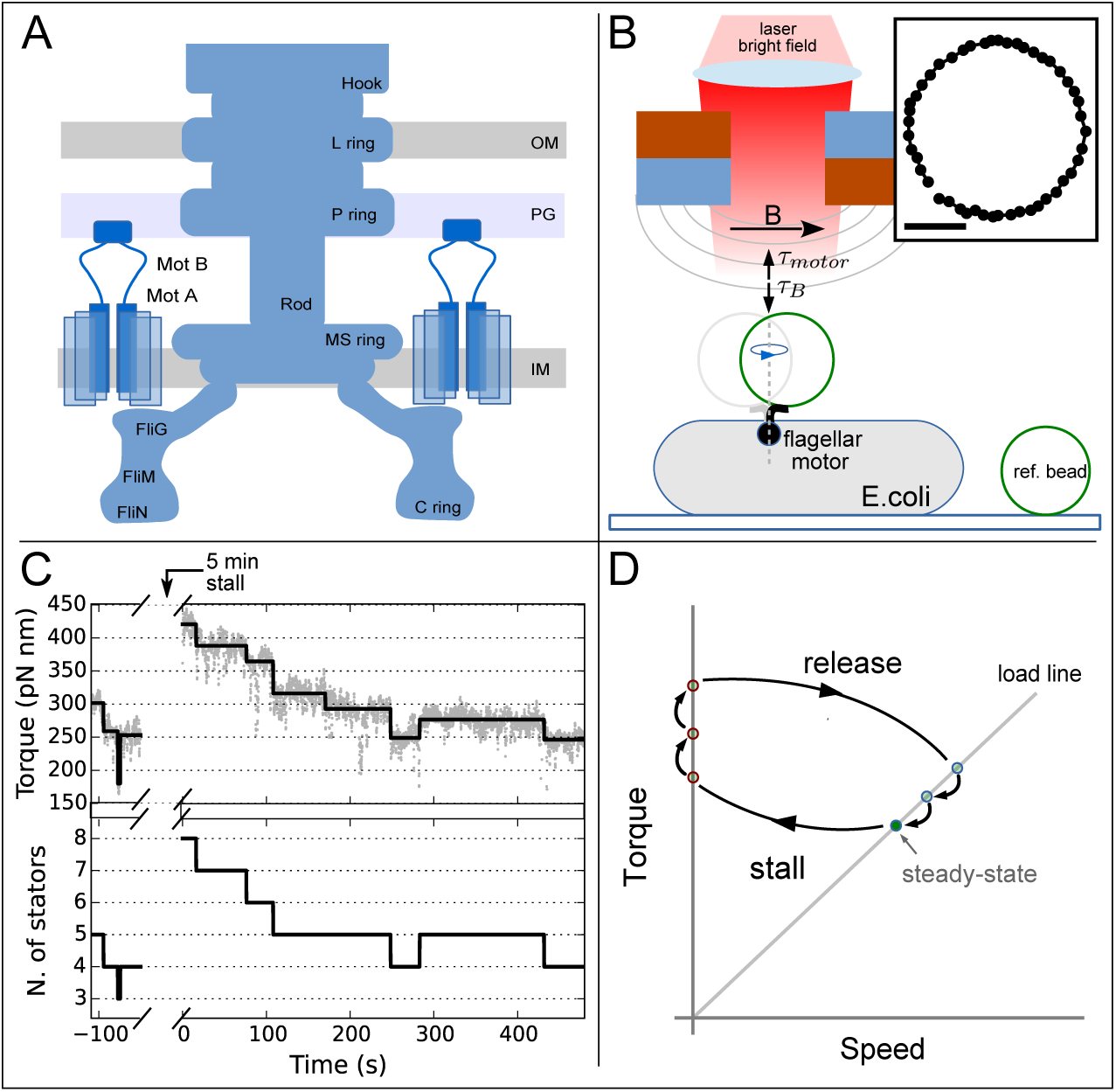
The experimental assay. (A) A schematic of the BFM showing the membrane embedded rotor and the stator protein complexes (comprised of MotA and MotB). Stator units which bind to the peptidoglycan at the periphery of the rotor provide torque via an interaction with FliG. (OM, PG, IM: outer membrane, peptidoglycan, and inner membrane) (B) Experimental setup. Bacterial cells are immobilized onto a coverslip, and a rotating superparamagnetic bead attached to the hook of a BFM is imaged and tracked. Drift correction is achieved by imaging a reference bead stuck to the cover slip in the same field of view. Two permanent magnets (aligned anti-parallel) create a magnetic field parallel to the cover slip. The magnitude of the magnetic field is modified by controlling the distance between the magnets and the sample. When sufficiently strong, the magnetic field generates sufficient torque *τ*_*B*_ on the bead to stall the motor. Inset: tracked positions of 1*μ*m bead rotating one turn (scale bar: 100 nm). (C) One experimental trace. Motor torque (top, gray points) is measured before stall by the magnets (*t <* 0), then again immediately upon release (*t >* 0). The output of the step detection algorithm (black line) is used to determine stator stoichiometry (bottom, see Materials and Methods). (D) The experimental assay shown in the torque speed plane. Starting from steady-state rotation (filled circle lying on the bead’s load line, i.e. the line *τ* = *γω*, where *τ, γ, ω* are the torque, the viscous drag coefficient, and the angular speed, respectively), the magnetic torque is increased to stall the bead. The motor is held stalled (*γ* = *∞, ω* = 0) for 5 min, during which time stator units may be recruited, increasing the motor torque (points outlined in red mark transitions inferred, not observed). The magnets are then moved far from the sample and the motor returns to the original load line, often with increased torque due to stator units incorporated during stall. The system eventually relaxes to the initial steady-state point as units dissociate.

Several different studies have revealed the continuous exchange of various BFM molecular constituents,^15–17^ demonstrating that a static model for the BFM structure is not adequate. A prime example, and in contrast to macroscopic rotary motors, the stator units of the BFM are dynamic; while each bound stator acts upon the rotor independently,^18, 19^ their stoichiometry in the motor varies. Once anchored around the rotor, stator units dynamically turnover, eventually diffusing away in the inner membrane.^17^ Each additional recruited stator increases the total torque and thus the measurable rotational speed of the viscous load attached to the rotor.^18, 20^ Up to eleven stator units have been observed to engage in an individual motor in *E.coli*, ^17, 21–23^ while either 8 or 16 have been indirectly inferred.^20, 24^ In these experiments, the number of active stator units has been modified via their expression level, and was found dependent on the cellular ion motive force.^25, 26^

Recently, novel observations have revealed that the stator units are also mechanosensors.^27–29^ There are a variety of mechano-sensitive membrane protein complexes which exist in all three kingdoms of life.^30^ While these complexes vary widely in their structure, function, and sensitivity, they share one key feature: the conformational state, and thus the function, of the protein is directly dependent upon mechanical stress, mediated by the surrounding cell membrane.

Two recent works have shown that stator recruitment in the BFM depends on the viscous load placed upon the motor.^27, 28^ The property of mechano-sensing (likely relevant for the cell to overcome local inhomogeneities and obstacles) has important consequences for the interpretation of previous data and ultimately for successfully modeling the torque generation of the BFM. It implies, in fact, that previously measured torque-speed relationships^18, 31–33^ are likely comprised of motors with a dynamically changing numbers of stator units. Theoretical models must now take this novel fact into consideration.^19^ Here, to better elucidate the molecular mechanism responsible for the mechano-sensitivity of the BFM, we investigate and characterize single motors by novel means. Using an external magnetic field and magnetic micro-beads of different sizes bound to the hook of individual *E. coli* motors, we rapidly manipulate the load experienced by the motor by reversibly stalling its rotation. Furthermore, we have developed a novel algorithm, which incorporates a recently developed step-detection algorithm,^34, 35^ to estimate the number of active stator units from high-resolution torque measurements of single motors. The external load manipulation directly probes the mechano-sensitivity of the BFM: we stimulate stator binding during the period of stall and observe and quantify stator unbinding after release. We perform these experiments for various viscous loads, each of which imposes a different initial stator occupancy. This allows us to statistically characterize the kinetics of stator stoichiometry both in steady-state conditions and following a rapid change in external load. Our analysis shows that the rate of stator attachment is independent of viscous load, while disengagement is load dependent, decreasing with increasing single stator torque. It is this effect that causes the observed mechano-sensitivity: the higher the torque produced by a single stator, and by symmetry, the higher the tension between the stator and its membrane anchoring point, the longer its lifetime in the motor complex. We thus propose that a catch-bond mechanism (a bond counter-intuitively strengthened, instead of weakened, by force)^36–39^ is at the heart of BFM mechano-sensitity, dynamically remodeling stator stoichiometry against changes in external resistance to rotation.

## Materials and Methods

### Bacteria and experimental configuration

We employ *E. coli* strain MTB32, a derivative of RP437. In MTB32, *FlgE*, the gene expressing the hook, has been replaced with an Avidity tagged peptide fused *FlgE* for conjugation of biotin.^40^ Additionally, we genetically deleted *CheY*, the chemotactic response regulator, via Red/ET recombination (Gene Bridges). Frozen aliquots of cells (100 *μ*l, grown to saturation and stored in 25% glycerol at −80^*°*^C) were grown in Terrific Broth (Sigma-Aldrich) at 33^*°*^C for 5 hours, shaking at 200 rpm. The final OD_600_ was 0.5–0.6. Cells were immobilized to a poly-Llysine (Sigma-Aldrich, P4707) coated coverslip (Menzel-Gläser) in custom made flow slides. Streptavidin superparamagnetic beads (1.36 *μ* m, Sigma-Aldrich, 543 nm or 302 nm, Adamtech) were washed in PBS (Sigma-Aldrich), resuspended in motility buffer (10 mM potassium phosphate, 0.1 mM EDTA, 10 mM lactic acid, pH 7.0), then allowed to spontaneously attach to the biotinylated hooks. Experiments were performed in motility buffer at 22^*°*^C. The sample was illuminated using a 660 nm laser diode (Onset Electro-Optics, HL6545MG) on a custom-built inverted microscope. The hologram of rotating beads was imaged via a 100x 1.45NA objective (Nikon) onto a CMOS camera (Optronics CL600x2/M) at 1000 Hz. The *x, y* position of the rotating bead was determined using cross-correlation analysis of the bead image.^41, 42^ Beads attached directly to the poly-L-lysine were used as fiducial markers for drift correction. An ellipse was fit to the drift-corrected *x, y* positions to yield the angular positions. Ellipses were transformed to circles under the assumption that the observed elliptical trajectories are a projection of a tilted circle. An example trajectory is shown in the inset of Fig. 1B. The resulting angle and speed traces were median filtered using a window of 0.5 s. All analysis was performed with custom LabView, Python, and MATLAB scripts. Two magnets (aligned anti-parallel, NdFeB, Supermagnete)^43^ were mounted above the sample plane onto a linear motor (Standa) which controlled the distance between the magnets and the sample plane. Full movement of the motor is achieved in less than 3 s.

### Motor torque calculation and fitting

The rotational viscous drag coefficient of a bead bound to the hook of a functional BFM was calculated as

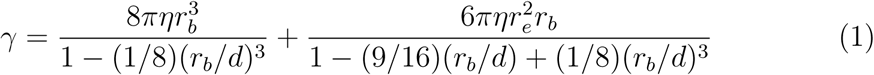

where *η* is the viscosity of the surrounding buffer, *r*_*b*_ is the radius of the bead, *r*_*e*_ is the measured radial distance to the bead’s axis of rotation, and *d* is the distance from the bead to the cell surface, estimated to be 5 nm in our experiments. In Eq. (1), the denominator corresponds to Faxen’s corrections.^44^ The torque which the motor applies upon the viscous load of the bead is *τ*_*motor*_ = *γω* where *ω* is the measured bead’s rotational velocity.

Torque traces were assumed to be noisy piecewise constant signals. A minimization of the *L*^1^-Potts functional was used to recover the underlying piecewise constant torque signal for both the pre-and post-stall motor torque traces. This was done via the PottsLab 0.42 toolbox in Matlab.^35, 45^ See the Supplementary Text for further details. An example torque trace and its fit is shown in Fig. 1C. This algorithm is ideal as it allows for the identification of discrete discontinuities, or steps, in the torque signal, while not requiring them to be of any specific size or the same size.

### Stator stoichiometry calculation and analysis

Stator stoichiometry was determined by preserving the discrete discontinuities from the step detection algorithm. As in previous works, the steps found in the torque traces are interpreted as indications of a change in stator number.^18, 20, 21, 24, 46^ For each individual motor, the average step size was used to determine the stator number as a function of time (see SI for more details and a test of both the step detection and stator stoichiometry determination algorithms).

Kernel density estimate distributions (Fig. 2A-C) were constructed with a Gaussian kernel (width of 1/2 the standard deviation of all fit torque steps in Fig. 2A, width of 1 stator in Fig. 2B-C). Average single-stator torque values were determined by performing a two-Gaussian fit to these distributions, where only the mean *m*_1_ and standard deviation *s*_1_ of the first Gaussian were free parameters, while the second Gaussian mean was set to *m*_2_ = 2*m*_1_ and the standard deviation was set to 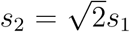. This considered large torque discontinuities as unresolved double steps, and allowed to better fit the long tail of the distribution. Single stator local force values (Fig. 3A-C) were calculated as the average single-stator torque divided by the radius of the rotor (23 nm^47^).

**Figure 2:**
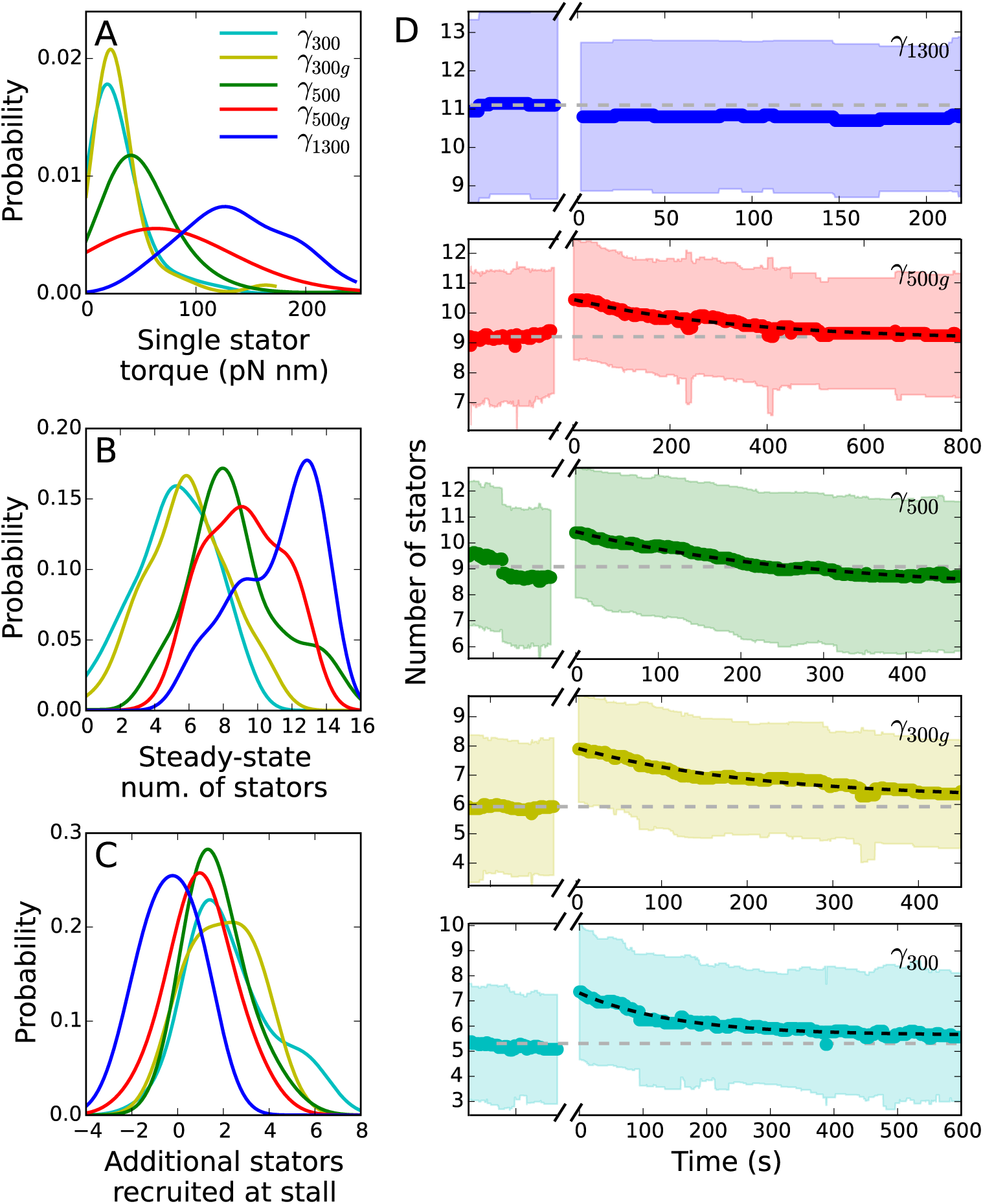
Stator stoichiometry before and after stall. (A-C) Kernel density estimates of the single-stator torque contribution, the steady-state stoichiometry, and the number of stator units recruited during stall, respectively, as a function of external viscous load. (D) Temporal evolution of stator stoichiometry of motors driving the different viscous loads (color-coded as in the other panels), before and after the forced stall. Steady-state rotation of the viscous load corresponds to time *t <* 0. Following a steady state measurement, the motor is stalled by the magnetic field for a period of 300 s (not shown, indicated by a break in the x axis). At *t* = 0 the motor is released from stall. The thick color-coded line is the average of several traces obtained with different motors, the colored region shows the corresponding standard deviation. The horizontal gray dashed line indicates the global average number of stator units measured for *t <* 0 at steady-state. The dark dashed line is the fit obtained from Eq. (4) for *t >* 0. Number of motors analyzed: 24 for *γ*_300_, 28 for *γ*_300*g*_, 40 for *γ*_500_, 30 for *γ*_500*g*_, 20 for *γ*_1300_.

**Figure 3:**
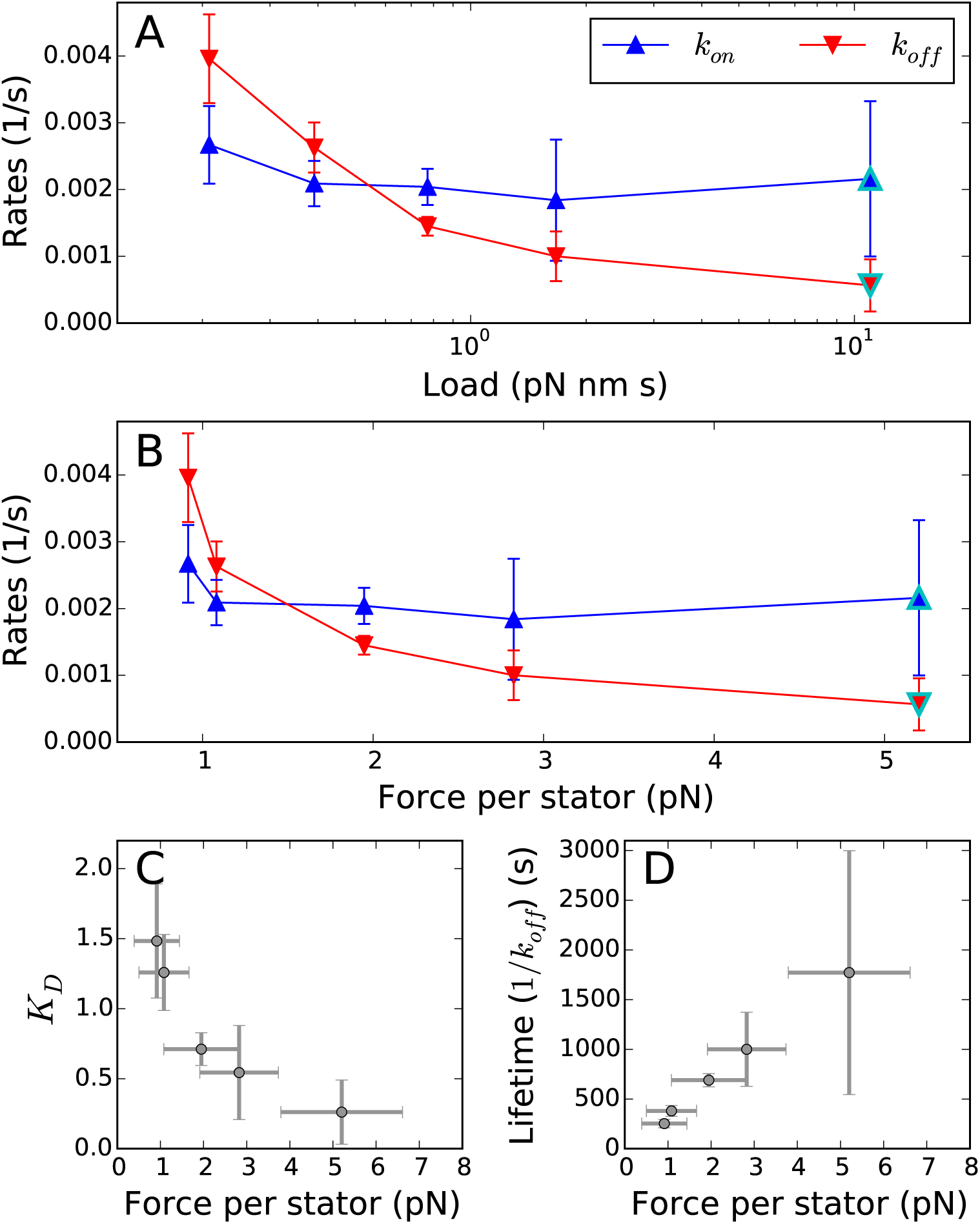
Stator kinetics. (A-B) The binding and unbinding rates of the stator units as a function of (A) external viscous load on the motor, and (B) single-stator force. All rates were calculated by fitting Eq. (4) to traces in Fig. 2D, with the exception of points outlines in cyan (see Supplementary Materials). (C-D) Dissociation constant, *K*_*D*_, and lifetime of an individual stator in the motor complex, respectively, as a function of the average local force applied by a single stator to the rotor (and by symmetry to the PG layer). Points and error bars give averages and standard deviations, respectively.

For each viscous load except *γ*_1300_ (in which stall had no statistical effect on the motor), Eq. (4) was fit to the average stoichiometry data after stall, with *k*_*in*_, *k*_*off*_, *N*_*ss*_, and *N*_*o*_ left as free parameters (least squares minimization, black dashed line for *t >* 0 in Fig. 2D). From the distributions of *N* at steady-state shown in Fig. 2B (and considering the uncertainty of these measurements), in this analysis we fix *N*_*max*_ = 14, which is compatible with previous estimates of 11-16.^17, 21–24^ However, our main conclusions do not critically depend on this choice. The errors of the fit parameters shown in Fig. 3A, were estimated by bootstrapping, using the standard deviation of the set of parameters that fit 100 random sub-samples (each containing 90% of the original set of traces) for each viscous load. To quantify the rates at *γ*_1300_, given the the relatively constant value of *k*_*on*_ for the smaller viscous loads, we made the assumption that *k*_*on*_(*γ*_1300_) is equal to the average of *k*_*on*_ for the smaller viscous loads. From the value of *N*_*ss*_(*γ*_1300_) we then obtained *k*_*off*_ (*γ*_1300_) from Eq. (3) (points in cyan in Fig. 3A-B)

In addition to the Hill-Langmuir adsorption model described by Eq. (4), we also considered a generalized reversible random sequential adsorption (RSA) model which incorporates a continuous binding ring around the rotor. As this model does not have an analytical solution. we determined stator binding and unbinding rates by using a Genetic Algorithm (Differential Evolution) to match simulated stator stoichiometry time trajectories to the average of the experimental trajectories. See SI for details.

## Results

### Torque Measurement and Load Manipulation

A non-switching strain of *E. coli* lacking flagellar filaments and containing an endogenously biotinylated hook was used for all experiments. Streptavidin coated superparamagnetic beads were attached to the hook of cells immobilized on a cover slip, and the rotation of the beads was observed via wide-field holographic microscopy^42^ (Fig. 1B). Tracking the position of the bead in time, we calculated the velocity and torque as described in Materials and Methods. Data were acquired for motors driving five different viscous loads (termed from high to low *γ*_1300_, *γ*_500*g*_, *γ*_500_, *γ*_300*g*_, *γ*_300_), which were obtained using beads of three different diameters and two buffer solutions of different viscosity, as indicated in Table 1. Two permanent magnets were mounted above the sample plane in a vertical orientation as shown in Fig. 1B. The magnets were attached to a fast motorized vertical translation stage which controlled the distance between the magnets and the sample, and thus the magnitude of the magnetic field at the sample plane. Both the BFM and the magnetic field exert a torque on the magnetic bead. For sufficiently large magnetic fields, the bead, and thus the motor, remain stalled in an equilibrium angular position where the magnetic torque and the motor torque cancel.^48^

**Table 1:** Angular drag coefficients used. Glycerol is listed in percent by volume. Bead diameter was measured by scanning electron microscopy.

| symbol | bead diam. | glycerol | drag (mean±st.dev.) |
| --- | --- | --- | --- |
| $\gamma_{1300}$ | $1364 \pm 98$ nm | 0 | $11.4 \pm 1.1$ pN nm s |
| $\gamma_{500g}$ | $543 \pm 148$ nm | 25% | $1.67 \pm 0.27$ pN nm s |
| $\gamma_{500}$ | $543 \pm 148$ nm | 0 | $0.77 \pm 0.20$ pN nm s |
| $\gamma_{300g}$ | $302 \pm 40$ nm | 25% | $0.39 \pm 0.12$ pN nm s |
| $\gamma_{300}$ | $302 \pm 40$ nm | 0 | $0.21 \pm 0.11$ pN nm s |

For a given bead size, the steady-state rotation of individual motors was measured under a negligible magnetic field, for 50 s to 300 s prior to manipulation. The magnets were then lowered to a distance such that the magnetic torque could permanently stall the motor rotation, and the motor was held at stall for 300 s. The magnets were then raised to the original height, the viscous load returned to that supplied by the bead in its viscous environment, and the rotation of the motor was recorded for at least another 5 min. Each movement of the magnets occurred within 3 s. The load felt by the BFM was therefore quickly and dynamically manipulated twice during the measurement of an individual motor; an exemplification of this procedure is shown in the torque-speed plane in Fig. 1D. The torque of individual motors was measured first at steady-state prior to stall and then again immediately after stall. An example torque trace for the viscous load *γ*_500_ is shown in Fig. 1C (top panel, see also Supplementary Materials Fig. S1 for a collection of individual traces at different viscous loads). An increase in torque during stall is visible, followed by a step-wise relaxation to a torque value close to the original steady-state value.

### Viscous Load Dependency of Stator Assembly Dynamics

Under the assumption that torque traces represent noisy constant signals demarcated by discrete discontinuities due to stator association or dissociation, we used a recently developed step detection algorithm^34, 35^ to fit the individual torque traces. Using this fit, a newly developed algorithm (see Methods and SI) was employed to calculate stator stoichiometry, extracting stator number as a function of time, *N* (*t*), for each individual trace. Contrary to previous works, this algorithm determines stator stoichiometry based upon the discrete discontinuities of the torque traces, not upon the absolute value of the torque; given the broad distributions of single-stator torque values (see Fig. 2A), this approach greatly reduces the error in stator stoichiometry estimation. Simulations (see Fig. S2) suggest that, given the average noise in our torque measurements, the algorithms used here are able to reconstruct the stator stoichiometry with an accuracy of 1.6 stator units, and that the shortest states in stator stoichiometry which can be reliably resolved last 3.5 s.

In Fig. 2 we show the statistical results of the analysis of the number of stator units *N* (*t*), on motors both at steady-state and after a 300 s period of stall. From the change in torque produced by resolved stator association and dissociation events, we quantified the distribution of torque produced by a single stator (Fig. 2A). This analysis shows that the torque generated by a single stator increases with increasing viscous load (and decreasing speed), matching theoretical models of stator behavior,^19, 49^ and confirming for MotAB stator units previous results based on sodium-driven PomAB stator units.^50^ Additionally, at steady state, we observed that the average stator occupancy is proportional to the viscous load applied (Fig. 2B and Fig. S3); this dependency is the fingerprint of the stator mechano-sensitivity.

In Fig. 2D, we show for each viscous load the average (colored line) and standard deviation (shaded region) of the number of stator units *N* (*t*) obtained from different motors, before and after stall. For all the viscous loads except for the highest (*γ*_1300_), the average stator number increases during stall. This is quantified in Fig. 2C: stalling a viscous load *γ*_1300_ for 300 s does not yield a relevant change in stator number (− 0.3 *±*1.1), while for the other viscous loads, mechano-sensing causes the recruitment of additional stator units during stall (1.0 *±*1.3 for *γ*_500*g*_, 1.7 *±* 1.1 for *γ*_500_, 2.0 *±* 1.4 for *γ*_300*g*_, 2.3 *±* 1.8 for *γ*_300_). After stall, as visible in Fig. 2D, *N* (*t*) decays back to the pre-stall, steady-state value within *∼*200 −300 s. This implies that, on average, for all the viscous loads except for the largest, additional stator units bind and engage with the BFM during the 300 s the motor is stalled. Within minutes after the magnetic field is removed and rotation resumes under the original viscous load, stator units dissociate and their average number returns to the previous steady-state value, which depends on the viscous load experienced during rotation. This behavior is not observed at the highest viscous load *γ*_1300_, which shows, on average, the same torque after stall as before, indicating that no statistically relevant change in stator number occurs during stall for this high viscous load.

### Modeling Stator Assembly Dynamics

In order to determine the stator binding and unbinding rates, a model of stator assembly is required. The simplest model of stator assembly kinetics, previously employed implicitly,^17, 25^ can be written as a Hill-Langmuir adsorption model.^51^ This model describes the rotor as surrounded by *N*_*max*_ independent and non-interacting binding sites. A diffusing stator can bind to an empty site with a constant rate *k*_*on*_, while a bound stator can disengage with a constant rate *k*_*off*_. The resulting average stator occupancy *N* (*t*) follows the dynamics

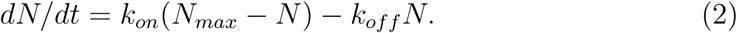

Here the concentration of unbound stator units is considered constant and unaffected by the binding and unbinding events. At steady state, *dN/dt* = 0, and the steady-state stator occupancy, *N*_*ss*_, is determined by

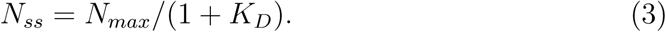

where *K*_*D*_ = *k*_*off*_ */k*_*on*_ is the dissociation constant. Under steady-state conditions, previous observations of stator turnover^17^ can be explained by the re-establishment of the steady-state number of stator units against fluctuations. This model is analogous to reversible RSA models;^52, 53^ in this case, the discrete lattice takes the form of a circle at the periphery of the rotor.

In line with previously published experiments demonstrating stator mechanosensing,^27, 28^ the viscous load-dependent distributions of *N* (*t*) that we measure under steady-state conditions, *N*_*ss*_, shown in Fig. 2B, imply that *K*_*D*_ depends on the applied viscous load *γ*, with a decreasing dissociation rate with increasing viscous load. Rapidly stalling a motor is equivalent to switching its viscous load to infinite (*γ→ ∞*), effectively shifting the steady-state towards *N*_*max*_. Accordingly, in our experiments, except for the viscous load *γ*_1300_, we observe a clear increase in the number of stator units after stall, indicating stator assembly upon an increase in applied load. A load-dependent *K*_*D*_ indicates either a load-dependence in *k*_*on*_, a load-dependence in *k*_*off*_, or a combination thereof.

In order to investigate this further, we analyzed the relaxation traces by comparing them with the analytical solution of Eq. (2), which predicts an exponential decay towards the steady-state occupancy *N*_*ss*_,

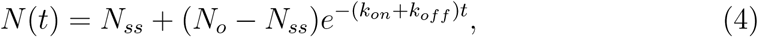

where *N*_*o*_ is the observed stator occupancy after the stall, starting the relaxation process (*t* = 0). The experimental mean traces for *N* (*t*) after stall, shown in Fig. 2D, are well fit by a single exponential; this simple model of stator binding kinetics is thus compelling, and it allows the calculation of the binding and unbinding rates from the experimental traces using Eq. (2).

The kinetic rates extracted at the different viscous loads are shown in Fig. 3A. It is evident that the differences in relaxation after stall are mainly due to a change in *k*_*off*_ with viscous load, while the values of *k*_*on*_ are relatively independent of viscous load. For the highest viscous load *γ*_1300_, we find that *N* does not change during stall and the exponential relaxation is absent (Fig. 2D), so the rates cannot be extracted. By contrast, this suggests that *K*_*D*_(*γ*_1300_) *∼ K*_*D*_(*γ* =*∞*), i.e. that the BFM mechano-sensitivity saturates for high viscous loads *γ* ⩾ *γ*_1300_, and that there is no dynamical difference between rotating such a high viscous load and being stalled. To quantify the rates at *γ*_1300_, given the the relatively constant value of *k*_*on*_ for the smaller viscous loads, we make the assumption that *k*_*on*_(*γ*_1300_) is equal to the average of *k*_*on*_ for the smaller viscous loads. From the value of *N*_*ss*_(*γ*_1300_) we can then obtain *k*_*off*_ (*γ*_1300_) from Eq. (3).

While the proposed Hill-Langmuir adsorption model is sufficient to explain and fit our experimental data, we note that there is no evidence for fixed binding sites at the periphery of the rotor in *E. coli*. Therefore, we also explored a more generalized reversible RSA model which is analogous to the classic ‘car parking problem’,^52–54^ where the stator binding is not restricted to a discrete number of binding sites but can occur continuously at any angular position on the ring (see Fig. S4A-B). In this model, a new stator cannot bind unless enough contiguous space is available in the ring (stator overlap is not allowed), which depends on the positions of the units currently bound. Hence, memory effects due to excluded-volume arise, affecting the stator occupancy dynamics, *N* (*t*). The details of this model are further discussed in SI. Numerical simulations of the model (see Methods and SI) exhibit similar relaxation dynamics to the experiments and the Hill-Langmuir model, and we conclude that both models can adequately fit our experimental observations. Strikingly, the extracted rates from fits of the experimental data are very similar to those of the Hill-Langmuir model (see Fig. S4), confirming the viscous load-independent binding probability combined with an unbinding probability that decreases for increasing viscous loads.

## Discussion

In this study, we have directly probed BFM mechano-sensing behavior, providing an extensive quantification of stator stoichiometry as a function of external viscous load, both at steady-state and immediately after a controlled change of load. By fitting measured stator kinetics to a reversible random sequential adsorption model, we provide the first measurement of stator association and dissociation rates as a function of external viscous load. We find that, while the rate of stator association is independent of viscous load, the rate of stator dissociation is loa dependent, and it is this property that begets mechano-sensitivity in the BFM.

Fig. 2A shows our measurements of *N*_*ss*_ as a function of viscous load, confirming previous results^27, 28^ that steady-state stator number is proportional to viscous load. The saturation curve of this relationship (see Fig. S3) is consistent with previous work which measured motor fluorescence as a function of viscous load (where the fluorescence signal of stator units fused to a fluorescent protein were a direct proxy of stator number).^28^ While this work provided evidence for a viscous load dependent *K*_*D*_, it was unable to identify if this dependence was governed by *k*_*on*_, *k*_*off*_, or both. A previous measurement of *k*_*off*_, ^17^ performed on immobilized cells where motors were presumably stalled via the attachment of flagella to the coverslip, reports a value which is two orders of magnitude faster than our measured *k*_*off*_ (*γ*_1300_). However, this study was performed with stator units fused to a fluorescent protein, which can cause different behaviors from their wild type counterparts.^17, 55^ The current study is the first, to our knowledge, to report on the dynamics of wild type stator units in otherwise unperturbed motors. Interestingly, our quantification of the viscous load dependence of stator association and dissociation rates has an important consequence for the molecular mechanism responsible for stator mechano-sensitivity in the BFM. We have seen that an increase in external viscous load translates into a higher torque, generated by each single stator on the rotor (as shown in Fig. 2A). Considering the average single-stator torque and the radius of the rotor (23 nm^47^), we quantify the mean local force applied by each single stator onto the rotor (found in the range of few pN, Fig. 3B). We note that, by reaction, this is also the force with which the stator simultaneously pulls upon and stretches its connection to the PG. In Fig. 3C and D, we show the measured dissociation constant *K*_*D*_ and lifetime (1*/k*_*off*_) of the stator as a function of this average force, finding that the lifetime of the stator increases with applied force. This counterintuitive relationship is the canonical fingerprint of catch-bond behavior.^36–39^ Whilst the lifetime of a conventional slip-bond decreases if tension is applied across it, a catch-bond produces a maximum of the lifetime at a non-zero force. A hook under tension and the Chinese finger-trap are two macroscopic analogies of this non-trivial molecular mechanism. We therefore conclude that the kinetic rates we measure suggest the existence of a catch-bond in the anchoring region of the stator on the peptidoglycan. We predict that if a force greater than the maximum force generated by a single stator could be applied to the anchor of the stator to the peptidoglycan, the catch-bond behavior will eventually transform into slip-bond behavior, as observed in other biological catch-bonds.^38^

The interface between the PG and the MotB PG-anchoring domain, located at the C-terminal of MotB (MotB_*C*_), is where the force can have an impact on the bonds relevant for the lifetime of the stator around the rotor. While our data indicates a catch-bond mechanism, it cannot discriminate any structural detail. However, interestingly, we note that it has been reported that the putative key PG-binding residues in the structure of the PG-binding dimer of MotB are buried and not readily accessible,^56^ and that a substantial structural flexibility of the domain is considered responsible to mask and unmask them.^57^ Recently, a conformational change upon binding has also been hypothesized in the PG-associated C-terminal of the closely related Outer Membrane protein OmpA.^58^ These facts suggest a possible catch-bond mechanism in which tension across the PG-MotB_*C*_ interface can promote the conformational rearrangements which lead to further exposure of binding residues to the PG, increasing the strength of the bond and the lifetime of the stator within the BFM complex (as sketched in Fig. 4).

**Figure 4:**
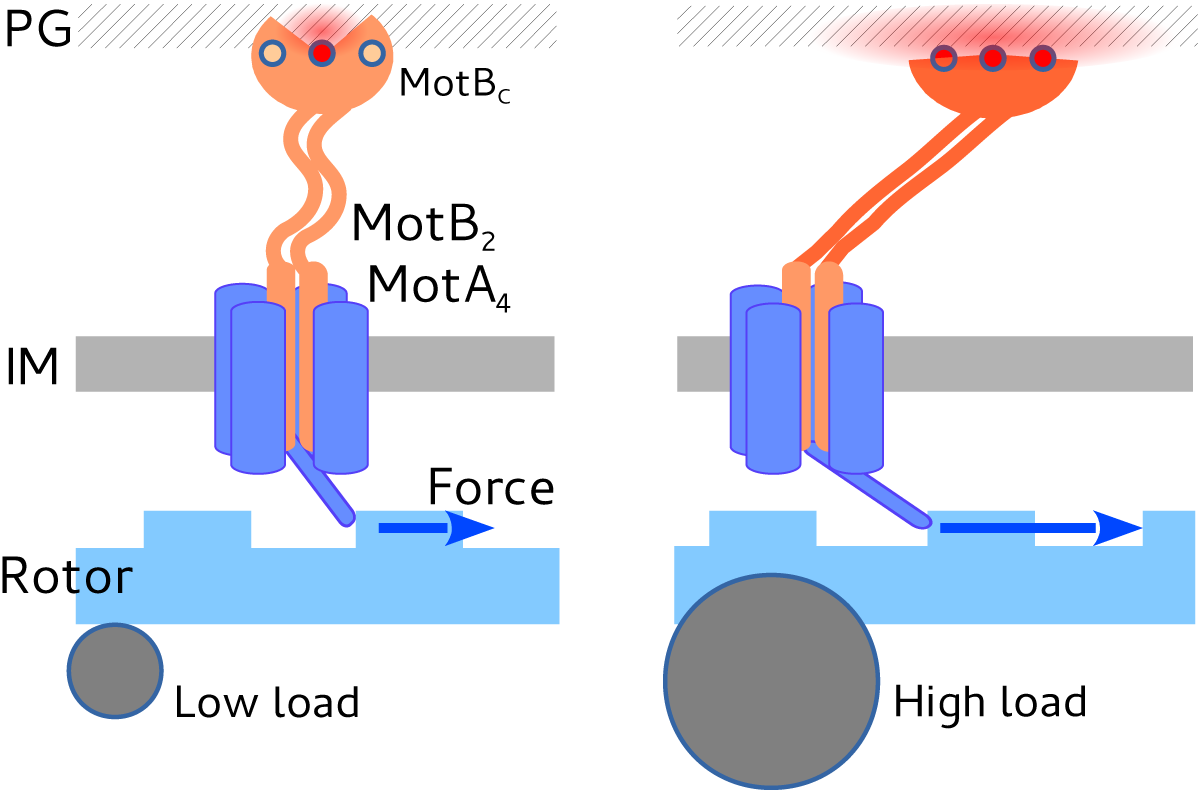
Cartoon of a proposed possible mechanism which increases the lifetime of each stator around the rotor when rotating a high viscous load (right) with respect to a low viscous load (left). The average force produced by the stator (blue arrow), by reaction, stretches the stator anchoring point at the PG, inducing conformational changes that increase the bond strength and lifetime.^65^ The average force is higher when a larger viscous load is rotated, consistent with previously published torque-speed curves, and as shown in Fig. 2A. (PG, IM: peptidoglycan and inner membrane).

As single molecule spectroscopy techniques continue to develop, the prevalence of experimental data demonstrating biological catch-bonds grows. Catch bonds have already shown to play an important role for two other molecular motors, myosin^59, 60^ and dynein.^61–64^ Here, we suggest that the mechano-sensitivity of the BFM may also be explained by a catch-bond mechanism within the stator. This feature potentially allows the cell to replace damaged stator units, adapt to the prevailing environmental viscosity, avoid wasting energy during flagellar growth, and may also play a role in behaviors which require surface-sensing, such as swarming motility and biofilm formation. We anticipate that future force-spectroscopy measurements, detailed structural studies, and molecular dynamic simulations will shed further light on the role of force in the anchoring of the stator units to the PG.

## Acknowledgments

We thank H.J.E. Beaumont and D. Dulin for constructive comments on this manuscript. We thank D. Chamousset for help modifying the bacterial strain. We thank M. Storath and A. Weinmann for *L*^1^ Potts Model step detection algorithms and support in implementation. CBS is a member of the France-BioImaging (FBI) and the French Infrastructure for Integrated Structural Biology (FRISBI), 2 national infrastructures supported by the French National Research Agency (ANR-10-INBS-04-01 and ANR-10-INBS-05, respectively). RPC acknowledges support from the Wellcome Trust (grant reference WT098325MA). ALN, EG, and FP acknowledge funding from the European Research Council under the European Union’s Seventh Framework Programme (FP/2007-2013)/ERC Grant Agreement no. 306475.

### Author Contributions

ALN and FP designed the experiments; ALN and EG performed the experiments; ALN, RPC, AB and FP analyzed the data; RPC, JAN, RMB, ALN, AB, and FP evaluated the kinetic models, ALN, RMB, and FP prepared the manuscript.

